# Connectome-based individual prediction of cognitive behaviors via the graph propagation network reveals directed brain network topology

**DOI:** 10.1101/2021.02.22.432377

**Authors:** Dongya Wu, Xin Li, Jun Feng

## Abstract

The brain connectome supports the information flow underlying human cognitions and should reflect the individual variability in human cognitive behaviors. Various studies have utilized the brain connectome to predict individual differences in human behaviors. However, traditional studies viewed the brain connectome feature as a vector of one dimension, a method which neglects topological structures of the brain connectome. To utilize topological properties of the brain connectome, we proposed that graph neural network which combines graph theory and neural network can be adopted. Different from previous node-driven graph neural networks that parameterize on the node feature transformation, we designed an edge-driven graph neural network named graph propagation network that parameterizes on the information propagation within the brain connectome. We compared various models in predicting the individual total cognition based on the resting-state functional connectome. The edge-driven graph propagation network showed the highest prediction accuracy and outperformed the node-driven graph neural network and traditional partial least square regression. The graph propagation network also revealed a directed network topology encoding the information flow, indicating that the high-level association cortices are responsible for the information integration underlying the total cognition. These results suggest that the edge-driven graph propagation network can better explore the topological structure of the brain connectome and can serve as a new method to associate the brain connectome and human behaviors.

## Introduction

Brain connectivity network provides pathways for transferring information between brain regions and serves as the basis of human cognition at a network level. Hence, individual differences in human cognitive behaviors should be reflected in the brain connectivity network. Investigating the brain connectivity network basis of these individual differences facilitates the understanding of brain network information processing mechanism that supports human cognitions and may benefit the developments of individualized medicine.

Numerous researchers have investigated the relationship between brain connectivity networks and individual differences in human cognitive behaviors. Smith et al. (2015) revealed a positive-negative axis linking human behavioral measures to a specific pattern of brain functional connectivity. Researchers have further adopted resting-state functional connectivity to predict human intelligence (Finn et al., 2015), sustained attention (Rosenberg et al., 2016) and creative ability (Beaty et al., 2018), or adopted functional connectivity derived from different task states to predict cognitive behaviors (Greene et al., 2018), and a connectome-based predictive modeling framework has been built to predict individual behaviors from brain connectivity (Shen et al., 2017). Zimmermann et al. (2018) also compared the unique and overlapping mapping of functional and structural connectivity on cognitive behaviors.

However, most current researches that investigate the relationship between brain networks and human behaviors treat the brain network features as a vector of one dimension, a method which neglects topological properties of the graph-structured brain network. To utilize these topological properties, one should treat the brain network as a graph instead of a vector, with connections between brain regions represented as edge features and regional properties represented as node features. Numerous studies have used graph theory to analyze the graph-represented brain network (Bullmore and Sporns, 2009), for instance, calculating the node degrees and central brain regions (van den Heuvel and Sporns, 2011), analyzing the small-world properties (Sporns and Zwi, 2004) and modularity properties of brain networks (Bullmore and Sporns, 2012), investigating the high order proximity properties of brain regions (Becker et al., 2018), or extracting various other graph metrics such as clustering, transitivity, integration (Kazeminejad and Sotero, 2018) and so on. However, the graph theory method needs to extract the graph features subjectively and cannot analyze the graph-structured data in an end-to-end fashion.

In view of the excellent performance achieved by the neural network and deep learning in extracting data features automatically (LeCun et al., 2015), it is promising to apply the neural network method to the graph-structured brain network to automatically learn the relationship between brain networks and human behaviors in an end-to-end fashion. The combination of neural network and graph-structured data is known as graph neural network (GNN). GNN has achieved rapid developments in recent years (Defferrard et al., 2016; Kipf and Welling, 2017; Veličković et al., 2018; Gong and Cheng, 2019; Xu et al., 2019) and obtained state-of-the-art results in many areas, such as the citation network, social network, and protein network, but the applications of GNN to the brain network are still very limited to date (Ktena et al., 2018; Parisot et al., 2018; Yang et al., 2019; Kim and Ye, 2020). In addition, these applications are based on the node-driven GNNs with parameters concentrating on the transformation of node features, a strategy that is not specialized for utilizing topological properties of the brain connectome. Edge-driven GNN as used by Wu et al. (2021) is needed to explore topological structures of the brain connectome.

To apply the GNN to the brain network analysis in a better way, we proposed an edge-driven GNN model named graph propagation network (GPN) (Fig. 1). The regional properties propagate within the brain network via connections between neighboring regions to form the representation of human behaviors. The relationship between brain networks and human behaviors is encoded in the propagation coefficients that are automatically learned by the error backpropagation. In this work, we utilized resting-state functional magnetic resonance imaging (rfMRI) data from the human connectome project (HCP) to construct the brain functional connectome. Different structures of the GPN were explored to investigate the connectome-based individual prediction of the total cognition. We further compared the GPN with the node-driven GNNs and partial least square regression (PLSR) that is widely adopted by traditional studies to demonstrate the priority of the GPN. Finally, we adopted the saliency map to investigate topological properties of the brain connectome underlying the total cognition.

**Figure 1.**
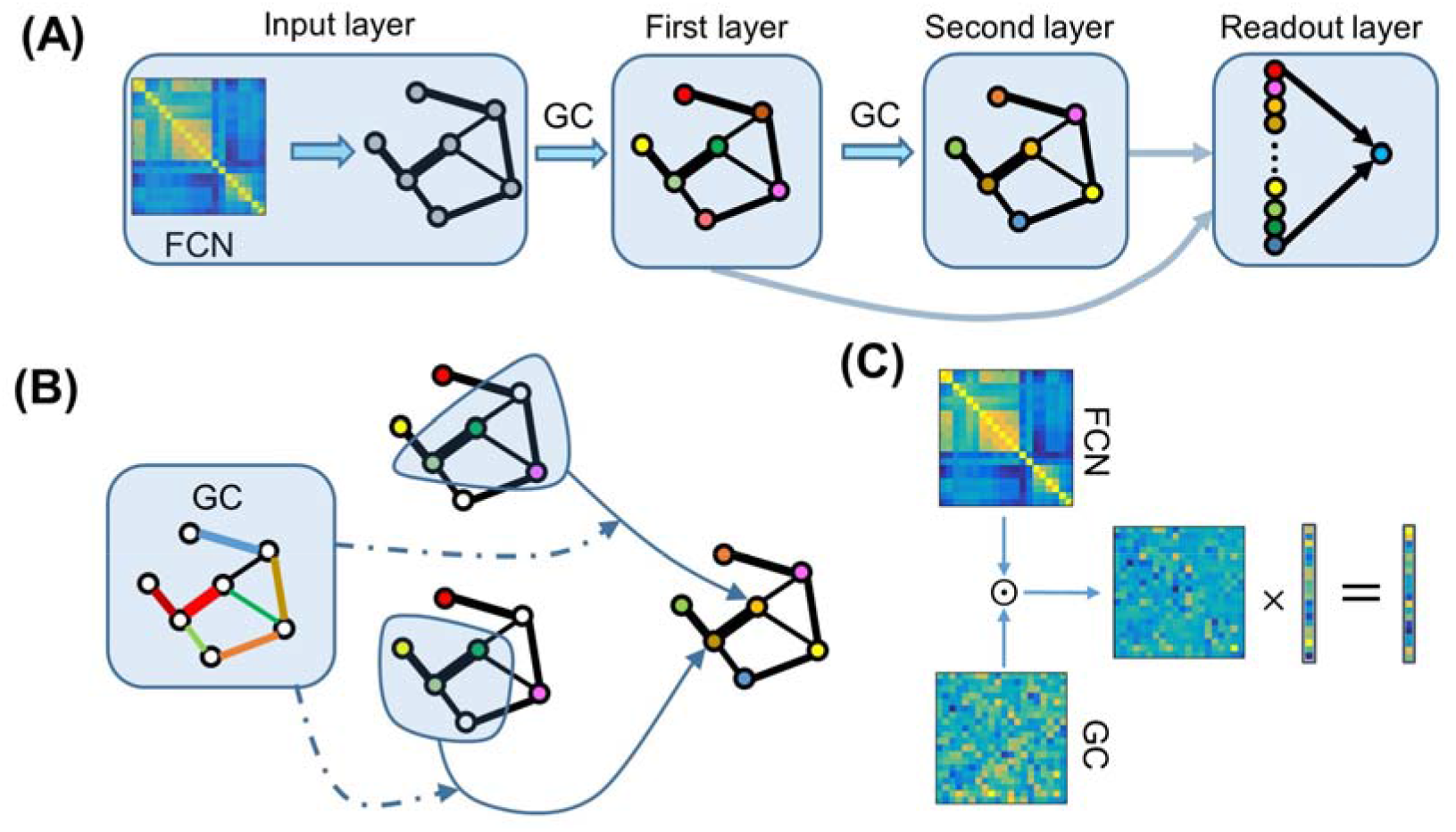
Schematic illustration of the GPN. (A) The graph-structured individual functional connectivity network serves as the input of the GPN. The node values are initialized as all ones. The graph convolution (GC) is the central computation of the GPN. The initial node values update through consecutive layers under the graph convolution. The node values of each layer are concatenated and the readout layer transforms this final graph representation to the individual prediction via a linear regression. (B) Each node updates its value by integrating the information from its neighbors. The graph convolution indicates the extent to which that each connectivity pathway participates in the information propagation. (C) The graph convolution can be realized by the matrix multiplication. The graph convolution parameters exert on the functional connectivity network via element-wise multiplication. The resulting matrix multiplies the vector that represents node value to update the node information.

## Materials and Methods

### HCP imaging and behavioral data

We adopted the rfMRI and behavioral data in the minimally pre-processed data provided by the HCP S1200 release (Glasser et al., 2013). We selected 997 subjects that have the rfMRI and behavioral acquisitions.

The rfMRI data were projected onto the 2 mm standard CIFTI grayordinates space, with the multimodal surface matching algorithm based on areal features used for accurate inter-subject registration (Robinson et al., 2014). Parameters acquisition and processing of the rfMRI data are described in Smith et al. (2013). Briefly, the rfMRI scans were acquired at 2 mm isotropic resolution, with a fast TR sampling rate at 0.72s using multiband pulse sequences (Ugurbil et al., 2013). Each subject had four 15-minute rfMRI runs, with a total of 1,200 time points per run. The rfMRI data were further processed by the ICA-FIX algorithm to remove the nuisances (Griffanti et al., 2014; Salimi-Khorshidi et al., 2014).

The behavioral data are described in Barch et al. (2013) and we adopted behaviors from the cognitive domain. As many cognitive measures are correlated, we used a single cognitive behavior “Cognition Total Composite Score” to measure the total cognition and further normalized this cognitive behavior to zero mean and unit variance. The “Cognition Total Composite Score” composes of both fluid and crystallized cognitive measures, with the fluid cognitive measures involving paradigms such as Flanker, Dimensional Change Card Sort, Picture Sequence Memory, List Sorting and Pattern Comparison, and the crystallized cognitive measures involving paradigms such as Picture Vocabulary and Reading Tests.

### Calculation of the functional connectome

We adopted the Human Connectome Project Multi-Modal Parcellation version 1.0 (HCP-MMP1.0) that contains 360 brain regions to define parcels of the brain (Glasser et al., 2016). Each subject’s four runs of rfMRI time series were demeaned, variance-normalized, and concatenated along the time axis successively. The time series of all vertices within each parcel were averaged. The averaged time series of each brain region were correlated with each other to form the functional connectome. We neither applied global signal regression on the rfMRI time series nor set any thresholds on the functional connectome, so as to avoid additional processing.

### Edge-driven graph propagation network

Fig. 1A shows the architecture of a 2-layer GPN. The GPN composes of three parts, i.e., the input layer, the middle layers (first and second layer in Fig. 1A), and the readout layer. In the input layer, the functional connectivity network constructed from the rfMRI time series is fed into the GPN as graph-structured data, with the node values initialized as ones. Each middle layer contains a graph convolution computation followed by an activation function. The graph convolution computation is central to the GPN and Fig. 1B shows examples of this computation. Each node integrates information from its neighbors to update its value, with the integration weighted by the strength of connections and graph convolution coefficients. The graph convolution computation can be realized by the matrix multiplication operation (Fig. 1C). In the readout layer, a linear model utilizes the node values of all layers to output the prediction of behavior.

Mathematically, a middle layer of the GPN can be represented in matrix form as follows:

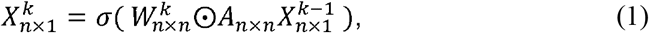

where *σ* is the activation function, *k* represents the *k*-th middle layer of the network, *n* represents the number of nodes in the GPN (*n* = 360 in this study), *A* represents the functional connectome, *W* indicates the extent to which each connection participates in the information propagation, and *X* represents the node value in the GPN. *W* exerts on *A* via the operation ⊙ which represents element-wise multiplication. The initialized node value in the *k*-th layer is a vector of ones: 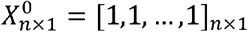. The node value in the *k*-th layer encodes the functional connectivity feature of *k* orders. To enrich the feature representation of the GPN, we concatenated the output of each layer to obtain the final representation as:

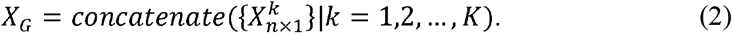

The readout layer utilizes a linear regression model to transform the final representation as:

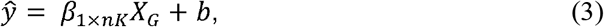

where *β* represents the transformation vector, *b* represents the bias term and *ŷ* is the prediction result.

### Node-driven graph neural network

We briefly described the node-driven GNN to make a comparison with the edge-driven GPN. A matrix representation of the node-driven GNN is as follows:

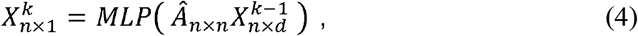

where MLP is the abbreviation of multi-layer perceptron that is used for node feature transformation, *d* is the dimension of node feature, and *X* represents the node feature that is the functional connectivity feature of each node. If we used a single-layer perceptron for feature transformation, model (4) can be represented as:

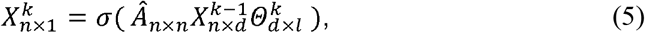

where *Θ* is the feature transformation parameter that transforms the node feature with *d* dimensions into a new feature with *l* dimensions, and *σ* is an activation function such as the rectified linear unit. If *Â* is the adjacency matrix *A_adj_* that is symmetrically normalized by the node degree, then model (5) corresponds to the graph convolutional network (Kipf and Welling, 2017). If *Â* = *A_adj_* + (1 + *ϵ*)*I*, where *I* is the identity matrix, then model (4) and model (5) correspond to the graph isomorphism network (GIN) (Xu et al., 2019). We also concatenated the node features of all layers and utilized linear regression to obtain the graph prediction output. Comparing model (2) with model (5), one sees that parameters of the edge-driven GPN concentrate on the propagation coefficient *W* and parameters of the node-driven GNN concentrate on the feature transformation *Θ*. These two parameterization strategies are not conflicted and one can also combine both parameterizations in one GNN. The combination of both edge-driven and node-driven strategies leads to a GNN as follows:

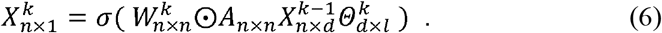

This combination method is also tested in this study.

### Training and testing procedures for the graph neural network

The dataset was randomly divided into a training set and a testing set with a ratio of nine to one. In the training stage, the mean squared error between the actual behavior and prediction was minimized to learn the parameters of the GPN. To ensure that the learning rate is not affected by the variance of each training batch, the mean squared error was normalized by the actual behavior’s variance of each training batch. In the testing stage, we used the Pearson correlation (*r*) to assess the prediction performance of the GPN. The value of *r* assesses the similarity between the actual and prediction value, and *r*^2^ approximates the proportion of individual behavioral differences that can be explained by the model. We performed this process 10 times with different random divisions of the dataset and used the mean Pearson correlation coefficient to assess the model.

### Saliency map that identifies the important connections

The propagation coefficient *W* cannot be directly used to indicate the extent to which each connectivity contributes to the individual prediction, as the GPN contains the batch normalization and the parameters of batch normalization also affects the information propagation. In order to calculate the importance of each connectivity feature, we adopted the saliency map method. The saliency map estimates each connectivity feature’s importance by calculating the extent to which the prediction result changes when each connectivity feature changes an infinitely small value. Mathematically, this amounts to calculating the gradient of each connectivity feature with respect to the prediction result. In practice, after we finished the training of the GPN, we calculated the gradient using each subject’s functional connectivity feature, and the final saliency map was calculated as the mean gradient across all subjects and 10 repetitions.

### Implementation details of the graph neural network

We implemented the GPN and GIN with PyTorch. The propagation coefficients in the GPN were initialized by the Xavier normal distribution with a gain of 0.1. The model was trained via the stochastic gradient descent optimizer with a Nesterov momentum of 0.9 and the training batch size was 128. The initial learning rate was 0.0001 for the GPN and a 0.1 multiplicative factor of learning rate decay was set at 250 epochs (300 total training epochs). The initial learning rate was 0.001 for the GIN and a 0.1 multiplicative factor of learning rate decay was set at 50 epochs (100 total training epochs). We adopted the batch normalization technique before the activation function and adopted the dropout technique in each layer’s output with a default rate of 0.5. We also added Gaussian random noise to the functional connectome of each training batch. The variance of the Gaussian random noise was equal to the variance of each connectome feature across subjects. We trained the models on an NVIDIA GeForce GTX 1080 Ti graphic processing unit.

## Results

### Comparison between graph propagation networks with different structures

We focused on three aspects of the GPN’s structures, i.e., shared or unshared parameter strategy, depth of the network, and choice of the activation function. Different middle layers can have different graph propagation coefficients (*W*) or shared the same graph propagation coefficient. As the size of the propagation coefficient in each layer is 360×360, the total parameters of unshared strategy grow more rapidly as the network gets deeper but the shared strategy controls the size of total parameters. We tested the performance of GPNs with different parameter strategies as the depth of the network gets larger under different choices of activation functions. As shown in Table 1, even though the total parameters of unshared strategies are several times than those of shared strategies, the GPNs with unshared strategies did not exhibit serious overfitting problems and showed comparable or even better prediction performance than those with shared strategies. In addition, GPNs that adopt linear activation functions showed better prediction performance than those that adopt rectified linear activation functions. In contrast to previous neural networks that need nonlinear activation functions to make the whole model nonlinear, the GPN can still achieve nonlinear mapping when linear activation functions are used. This nonlinear nature of the GPN resides in the information propagation process through consecutive layers.

**Table 1.** Prediction performance of GPNs. The accuracy is represented by the mean Pearson correlation and the standard deviation across ten random divisions of the dataset.

| Model | Parameter strategy | Layers | Activation function | Accuracy(*r*) |
| --- | --- | --- | --- | --- |
| GPN | Unshared | 1 | Linear/Relu | 0.545±0.055/0.525±0.052 |
| GPN | Unshared | 2 | Linear/Relu | 0.553±0.054/0.542±0.054 |
| GPN | Unshared | 3 | Linear/Relu | 0.551±0.053/0.537±0.051 |
| GPN | Shared | 2 | Linear/Relu | 0.553±0.054/0.524±0.049 |
| GPN | Shared | 3 | Linear/Relu | 0.546±0.047/0.540±0.031 |

### Contrasting the graph propagation network with the baseline model

We selected the GIN as a representation of node-driven GNNs, as the GIN is powerful than other GNN variants (Xu et al., 2019). The parameter *ϵ* is set as 0 for simplicity. We used each brain region’s functional connectivity features as the input node features. We derived the adjacency matrix in the GIN by setting a threshold on the functional connectivity matrix, i.e., the connections that have absolute values larger than the threshold are conceived to be positive or negative adjacent. To make the parameter size of GPN and that of GIN comparable, we firstly adopted a single layer MLP for node feature transformation and the dimension of features was set as 360 in the GIN. In this model setting, the parameter size is 360×360 in each layer of the GIN. In addition, we tested a 2-layer MLP for node feature transformation and the dimension of features was set as 64. The parameter size is 360×64+64×64 in each layer under this model setting. In these GINs, we adopted rectified linear activation functions rather than linear functions, as nonlinear activation functions are necessary for MLP to make the transformation nonlinear. Table 2 recorded the prediction performance of GINs under different settings. Setting a larger threshold on the functional connectivity network leads to GINs that showed better prediction performance. Under various model settings, the GPNs outperforms GINs when the parameter size is comparable or when GINs adopted a 2-layer neural network for node feature transformation.

**Table 2.**
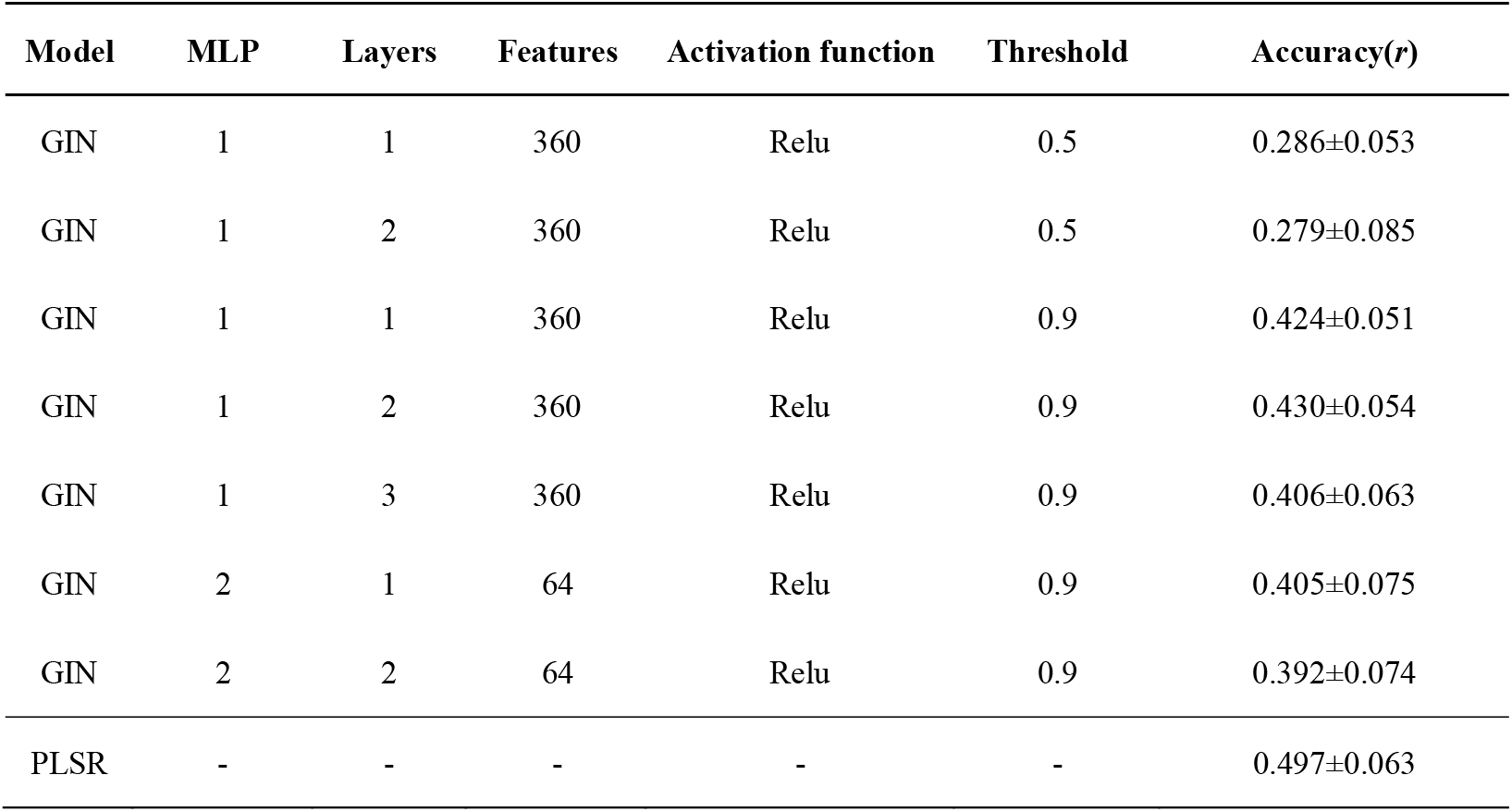
Prediction performance of baseline models.

| Model | MLP | Layers | Features | Activation function | Threshold | Accuracy(*r*) |
| --- | --- | --- | --- | --- | --- | --- |
| GIN | 1 | 1 | 360 | Relu | 0.5 | 0.286±0.053 |
| GIN | 1 | 2 | 360 | Relu | 0.5 | 0.279±0.085 |
| GIN | 1 | 1 | 360 | Relu | 0.9 | 0.424±0.051 |
| GIN | 1 | 2 | 360 | Relu | 0.9 | 0.430±0.054 |
| GIN | 1 | 3 | 360 | Relu | 0.9 | 0.406±0.063 |
| GIN | 2 | 1 | 64 | Relu | 0.9 | 0.405±0.075 |
| GIN | 2 | 2 | 64 | Relu | 0.9 | 0.392±0.074 |
| PLSR | - | - | - | - | - | 0.497±0.063 |

We also compared the GPN with the PLSR, as the PLSR performs well in the individual prediction based on the brain connectome (Yoo et al., 2018; Chen et al., 2019). As the functional connectivity network is symmetric, only the upper triangular part of the network is extracted and viewed as a vector for the PLSR. Divisions of the dataset for PLSR are the same as those for GNNs. We determined the number of PLSR components via ten-fold cross-validation on the training dataset and the model that achieved minimum cross-validation error was used for assessment on the testing dataset. We implemented the PLSR using the function “plsregress” in Matlab. Table 2 showed the prediction performance of the PLSR. All GPNs showed higher prediction accuracy than the PLSR but GINs showed lower prediction accuracy than the PLSR.

### Testing the combination of edge-driven and node-driven strategies

The edge-driven and node-driven GNNs differ in their parameterization strategies, with the edge-driven method focusing on the modeling of information propagation and the node-driven method focusing on the transformation of node features. We tested whether the combination of both strategies can improve the individual prediction of cognition based on the functional connectivity network (Table 3). Similar to the performance of GPNs, the combination strategy also showed better performance when adopting a linear activation function. The combination strategy showed higher prediction performance than GINs and showed lower prediction performance than GPNs. This result showed that adding graph propagation coefficient to the GINs can facilitate the utilization of graph’s topological structures and improve the individual prediction based on the brain connectome. However, adding the node feature transformation to GPNs reduces the performance of GPNs and the graph propagation parameterization is sufficient to utilize the brain network’s structure.

**Table 3.** Prediction performance of models with the combination strategy.

| Model | MLP | Layers | Features | Activation function | Parameter strategy | Accuracy(*r*) |
| --- | --- | --- | --- | --- | --- | --- |
| GPN+GIN | 1 | 1 | 360 | Relu | Unshared | 0.447±0.059 |
| GPN+GIN | 1 | 2 | 360 | Relu | Unshared | 0.438±0.063 |
| GPN+GIN | 1 | 1 | 360 | Linear | Unshared | 0.494±0.071 |
| GPN+GIN | 1 | 2 | 360 | Linear | Unshared | 0.475±0.057 |

### The directed network topology underlying the total cognition

We used the saliency map to visualize the functional connectivity network features of the 2-layer GPN (adopting unshared strategy and linear activation function) in Fig.2. As the functional connectivity network is symmetric, the feature pattern of traditional models such as the PLSR is also symmetric. However, the functional network features of the GPN showed an asymmetric pattern that derives from the asymmetric information propagation. Since the information propagation and integration are realized by the graph convolution in Fig. 1C, rows of the saliency matrix represent regions that integrate information from neighboring regions, and columns of the saliency matrix represent regions that send information out. Thus the saliency matrix revealed a directed network topology that encodes the information flow. One can identify each functional connection’s importance relative to the total cognition via the saliency matrix in Fig.2. We further calculated the node degree of each region as the summation of the absolute saliency matrix vertically or horizontally. The vertical node degree (on the left side of the saliency matrix) represents the importance of each region in integrating information from other regions, and the horizontal node degree (on the bottom side of the saliency matrix) represents the importance of each region in sending information out to other regions. The horizontal node degree showed a relatively homogeneous distribution pattern across cortices compared with the vertical node degree. The vertical node degree is plotted on the cortical surface and showed small values in the primary visual, auditory, and motor cortices, while showed large values in the high-level association cortices such as the dorsolateral and medial prefrontal, inferior frontal, and parietal cortices. This node degree distribution pattern indicates that the high-level association brain regions are responsible for the information integration supporting the total cognition.

**Figure 2.**
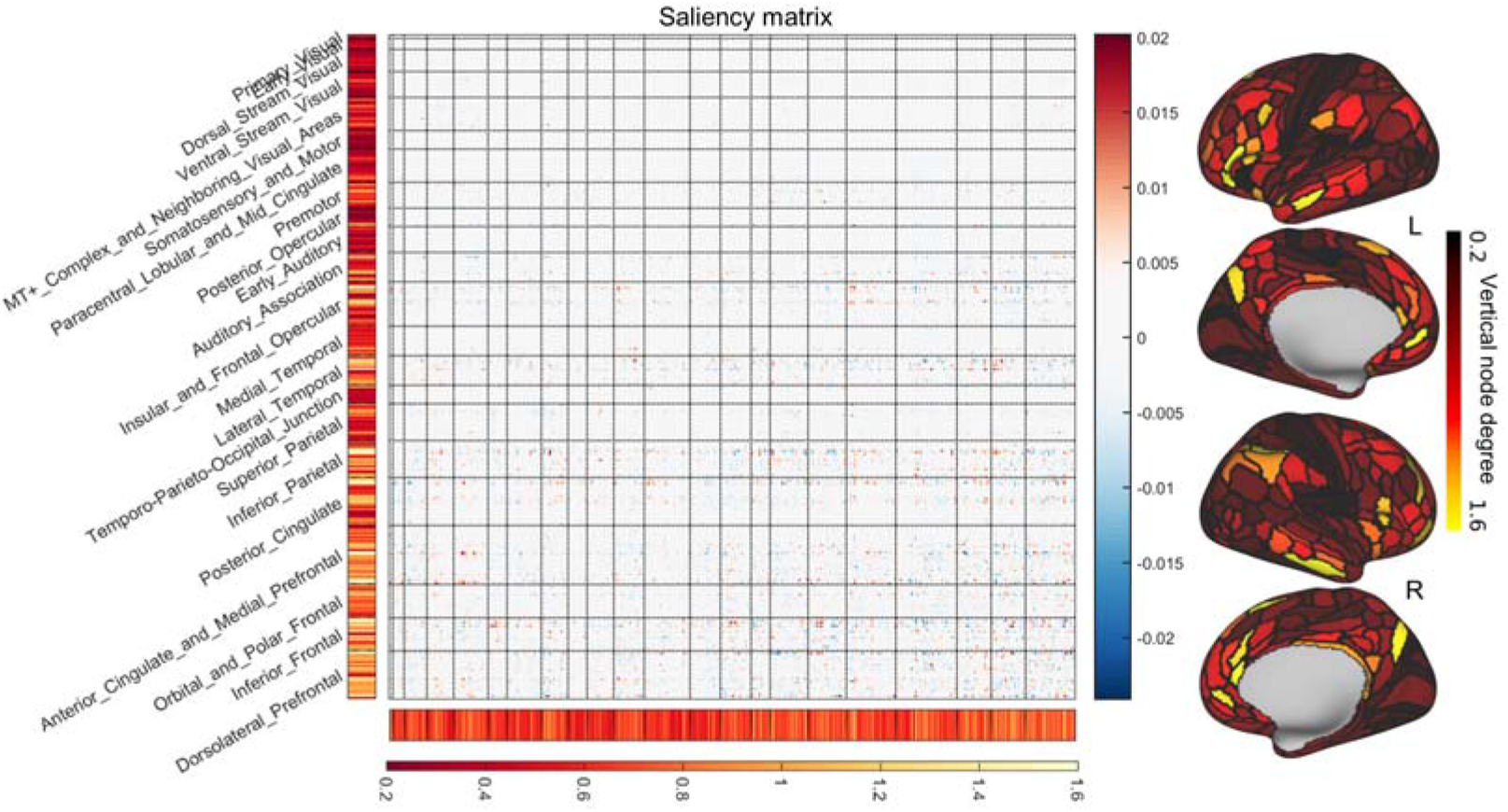
The saliency map of functional network features relative to the total cognition. The saliency map of functional network features is shown in the matrix. The saliency matrix value represents the gradient or importance of each connection with respect to the total cognition. The vertical and horizontal node degree of the saliency matrix are shown on the left side and bottom side of the matrix respectively. We grouped brain regions that belong to the same cortex and the cortical name follows the Supplementary Neuroanatomical Results for A Multi-modal Parcellation of Human Cerebral Cortex (Glasser et al., 2016). The vertical node degree is also plotted on the cortical surface to visualize the distribution pattern.

## Discussion

In the current study, we proposed an edge-driven GNN named GPN that can capture the individual association between the brain connectome and the human total cognition in a better way. Compared with either the traditional PLSR model that views the brain connectivity network as a vector and ignores the topological structure of the brain connectome or the node-driven GNN that adopts the feature transformation parameterization, the proposed GPN can model the information flow within the brain network and facilitate the use of the brain network’s topological structure. The edge-driven GPN exhibits higher performance than the PLSR and the node-driven GNN in predicting the individual total cognition based on the functional connectome and reveals a directed information flow supporting the total cognition. The GPN shows promising future in the field of individualized prediction based on the brain connectome.

He et al. (2020) showed that the graph convolutional neural network performs worse than traditional regression models, a study in which the authors adopted a node-driven GNN model (Kipf and Welling, 2017; Parisot et al., 2017; Parisot et al., 2018). Similarly, the results in our study also showed that another node-driven GIN performs worse than the traditional PLSR model. The previous node-driven GNNs and traditional regression models are not well adapted to process the graph-structured brain network data and cannot utilize the topological structure of the brain network. Since GNNs usually possess higher model complexities than traditional regression models, node-driven GNNs are prone to suffer overfitting problems and thus exhibited poor generalization abilities than simple regression models such as the PLSR. However, the edge-driven GPN tested in the current study exhibits superior performance compared with the traditional PLSR. This result indicates that GPNs which are specially designed for modeling information propagation in brain networks are more suitable for processing graph-structured brain network data.

We demonstrated that the edge-driven GPN is superior to the node-driven GNN in predicting individual total cognition from the resting-state functional connectivity network. This result suggests that viewing the functional connectome as the node feature may break the topological structure of the brain network, while the edge-driven GPN that can capture the information propagation within the brain network is better specialized for processing the graph-structured brain connectivity network. However, this does not mean that the edge-driven GPN is also better than the node-driven GNN in other conditions. In this study, we only utilized the functional connectome and did not adopt other brain node features, such as the cortical thickness, myelin, task activation and so on. If more node features are available to the GNN, the node-driven strategy that is better at node feature transformation may express a more powerful ability. Future works can incorporate more brain node features into the brain connectome and investigate whether the combination of edge-driven and node-driven strategies can facilitate the processing of multi-modality brain connectome.

Compared with the deep learning models that can contain over one hundred layers, the GPN adopted in this study contains only several layers. The reasons behind this shallow structure may be twofold. Since the brain network possesses the small-world property (Sporns and Zwi, 2004; Bassett and Bullmore, 2006), the minimum path length between any pair of brain regions is only several steps. Thus the functional information of each brain region can propagate to any other brain regions in the GPN within several steps. This short information propagation range within the brain may lead to the GPN with a shallow structure. On the other hand, the sample size of the data utilized in this study is only about one thousand, a number that is much smaller than that adopted in deep learning studies. As the depth of GPNs grows larger, the model complexity also gets larger, a condition in which adopting a small sample size is more prone to overfitting. Future works can adopt datasets with more samples, such as the biobank dataset (Alfaro-Almagro et al., 2018), to test whether adding more samples can improve the generalization performance.

The GPN revealed a directed information flow via the saliency map and the directed information flow further indicates that the high-level association cortices are responsible for the information integration that supports the total cognition. Even though traditional methods such as the PLSR can also reveal important connectivity features belonging to association cortices (Zimmermann et al., 2018; Wu and Li, 2019), the revealed features do not contain information about the direction of the information flow and researchers cannot identify whether association cortices send out or integrate information. The information flow within the brain network is essentially directed, but many functional or structural brain connectivity networks constructed via neuroimaging are undirected. Several methods such as Granger causality (Dhamala et al., 2008) and dynamic causal modeling (Friston et al., 2003) can estimate the directed connectivity based on neuroimaging time series. GPN estimates the directed information flow through a connectome-based individual prediction that links the brain network with human behaviors, a manner which differentiates GPN with those previous methods. A behavior context is necessary for deriving the directed information flow from the undirected brain connectivity network, as effective connectivity should be defined within a certain context (Park and Friston, 2013). Therefore, the revealed directed information flow is only a subset of the whole brain network and is relevant to the predicted cognitive behavior. The directed information flow revealed by the GPN also coincides with the view of neuroscience, as the high-level association cortices integrate functional information from primary cortices (Purves et al., 2001), which is important for supporting complex cognitions.

In conclusion, we proposed an edge-driven GPN that can greatly improve the individual prediction of the human total cognition from the brain functional connectome. The edge-driven GPN serves as a new kind of GNN relative to node-driven GNNs and focuses on the topological structure of the brain connectome. This advancement contributes to linking the individual differences in human cognitive behaviors to the brain connectome and suggests new perspectives on the information propagation mechanism within the brain connectome.

## Data availability

The HCP S1200 data release is publicly available online at https://www.humanconnectome.org/.

## Acknowledgments

Data were provided by the Human Connectome Project, WU-Minn Consortium (Principal Investigators: David Van Essen and Kamil Ugurbil; 1U54MH091657) funded by the 16 NIH Institutes and Centers that support the NIH Blueprint for Neuroscience Research.

## Author contributions

Wu, Li and Feng designed the research, Wu performed the data analysis, Wu, Li and Feng wrote and revised the paper.

